# Metachronal rowing provides robust propulsive performance across four orders of magnitude variation in Reynolds number

**DOI:** 10.1101/2024.11.18.624216

**Authors:** Mitchell P. Ford, Arvind Santhanakrishnan

**Affiliations:** School of Mechanical and Aerospace Engineering, Oklahoma State University, Stillwater, OK 74078, USA

**Keywords:** hydrodynamics, metachrony, swimming, bio-inspired robotics, biological propulsion, crustaceans

## Abstract

Metachronal rowing of multiple propulsors (paddles) is a swimming strategy used by numerous organisms across various phyla, with body sizes ranging from 0.01 mm to 100 mm. This size range corresponds to a huge variation in flow regimes characterized by Reynolds number (*Re*) ranging on the orders of 10^−2^ (viscosity dominated) to 10^4^ (inertially dominated). Though the rhythmic and coordinated stroking of paddles is conserved across species and developmental stages, the hydrodynamic scalability of metachronal rowing has not been examined across this broad *Re* range. We used a self-propelled metachronal paddling robot to examine how swimming performance changes across four orders of variation in *Re* (21 to 54,724) relevant to most aquatic crustaceans. We found that the Strouhal number (*St*), characterizing momentum transfer from paddles to the wake, was unchanging at *St* ≈ 0.26 for *Re >* 42 and within the reported *St* of various flying and swimming animals. Peak dimensionless strength (circulation) of paddle tip vortices linearly increased with *Re* and was mostly unaffected by changing fluid viscosity. Our findings show that the swimming performance of metachronal rowing is conserved across widely varying flow regimes, with dimensionless swimming speed scaling linearly with *Re* across the entire tested range.

## 1 Introduction

The metachronal rowing of discrete biological structures, such as cilia, parapodia, ctenes and limbs, is a fluid transport strategy used by aquatic organisms from a variety of different taxa. Metachronal rowing is used for several functions, including swimming [1–6], pumping [7], feeding [8, 9], and particle transport [10, 11]. When used for swimming, the propulsive elements (hereafter “paddles”) are rhythmically stroked in sequence starting from the posterior end of the animal and proceeding towards the anterior, resulting in a metachronal wave that travels in the same direction as the body motion. Organisms that employ metachronal rowing as a locomotion strategy have body lengths ranging on orders of magnitude from 10^−5^ m to 10^−1^ m [12]. This wide range of body sizes necessitates an equally wide range of hydrodynamic scales as quantified by the *Reynolds number* (*Re*). Reynolds number is defined as the ratio of inertial forces to viscous forces in a fluid flow, and for metachronal swimmers there are two different scales to consider: the appendage scale which is defined by the *paddle Reynolds number* (*Re*_L_) and determines the physics behind thrust production, and the body scale which is defined by the *swimming Reynolds number* (*Re*_B_) and determines the physics behind swimming resistance (hydrodynamic drag). A number of previous studies have given insight on the use of metachronal swimming by several species of crustaceans. At the propulsor scale, the motion of each paddle produces a region of negative pressure that generates propulsive thrust [13], and the flows produced by each paddle interact to enhance vorticity on the adjacent paddles [14, 15]. On the body scale the wakes of the individual paddles coalesce into a single near-continuous jet [8] which is detectable in a volume an order of magnitude larger than the body size [16, 17]. This large-scale wake contains hydrodynamic signals that can enable complex behaviors such as schooling [18] and mate tracking [19]. Despite the fact that paddling organisms are found across a large range of *Re*_L_ and flow regimes, it is unclear from these studies how this swimming strategy performs across size scales when the body planform and rowing kinematics are maintained.

Due to the differences in morphology and rowing kinematics across species, such single-species studies are limited in the insight that they can provide on mechanisms of force production across the wide range of hydrodynamic regimes experienced by metachronal swimmers. Numerical [20–22], bio-robotic [23, 24] and physical/robophysical [14, 25] models provide powerful alternatives to synthesize the unifying physical principles underlying metachronal swimming. Recent studies have applied these modeling approaches to investigate the functional roles of paddle kinematics and morphology, including the inter-appendage phase lag [14, 20, 21, 24, 26–30], appendage shape [31], inter-appendage spacing [27, 28], number of appendages [27, 32], and stroke amplitude [28, 32]. A common finding from some of these modeling studies is the formation of vortices at the tips of paddles during the power stroke (PS), which was originally observed in three-dimensional tomographic PIV measurements of hovering Antarctic krill at *Re*_L_∼ 10^2^ [18]. Using a self-propelling paddling robot at *Re*_L_∼ 10^2^, Ford and Santhanakrishnan [28] reported that tip vortex interactions in PS at biologically relevant dimensionless inter-paddle spacing resulted in increased horizontal momentum fluxes (surrogate of thrust) and swimming speed – relative to those generated by a paddling array with larger, non-biological dimensionless inter-paddle spacing frequently used in models. Destructive tip vortex interactions during metachronal recovery stroke (RS) have been reported to promote formation of shear layers on closely spaced paddles (as opposed to tip vortices in RS) that weaken flow generating “parasitic” drag (i.e., opposing thrust), thus augmenting average thrust over a stroke cycle [22, 28]. While these findings show the importance of paddle tip vortices in metachronal rowing performance, none of these studies have examined the effect of changing *Re*_L_ on tip vortex strength and interactions across the full range of *Re*_L_ typically experienced by adult crustaceans (10^1^ ≤ *Re*_L_ ≤ 10^4^).

Specific to Reynolds number scalability of metachronal rowing, Zhang et al. [21] conducted 2D numerical simulations of a tethered metachronal paddling model and found that increasing *Re*_L_ resulted in an increase in volumetric flux at *Re*_L_ ranging from 50 ≤ *Re*_L_ ≤ 800. Subsequently, Granzier-Nakajima et al. [27] found that volumetric flux increases with *Re*_L_ for 0.03 ≤*Re*_L_≤ 1, but decreases with *Re*_L_ for *Re*_L_ ≥ 1. Ford et al. [14] used a robophysical model to investigate the effects of *Re*_L_ and found that the direction of the wake, and therefore whether the propulsive force acts in the thrusting or lifting direction, varies with *Re*_L_ over a range from 32 ≤ *Re*_L_ ≤ 516. However, none of the existing studies have comprehensively examined swimming performance of metachronal rowing across varying *Re*_L_. In this study, we use a self-propelling robot to investigate how changing *Re*_L_ across four orders of magnitude from 10^1^ to 10^4^ affects the swimming performance and the wake generated by metachronal rowing, while maintaining the same paddle shape, spacing, stroke amplitude and inter-paddle phase lag.

## 2 Experimental Methods

### 2.1 Robotic model

A previously developed self-propelling metachronal paddling robot [24, 28, 29] was employed in this study to examine hydrodynamic scalability across four orders of magnitude of *Re*_L_. This model was validated [24] against published data [18] on hovering Antarctic krill (*E. superba*) using a geometrically scaled, krill-like body that was operated in a viscous fluid medium (water-glycerin mixture) in order to achieve dynamic similarity. In the current study, five flat plate-like paddles were used to serve as the robotic swimming paddles. Each paddle was actuated via a stepper motor to oscillate about its root following a sinusoidal motion profile. The motion of five stepper motors was controlled via a custom LabView program (LabView 2020, National Instruments, Austin, TX, USA). The motion control program read a matrix containing angular positions for each of the 5 motors prescribed at 10 ms increments. The same LabView program that was used to prescribe the sinusoidal motion profiles to the motors (Figure 1D) also sent electronic pulses that were used to synchronize the laser and high speed camera for data acquisition.

**Figure 1.**
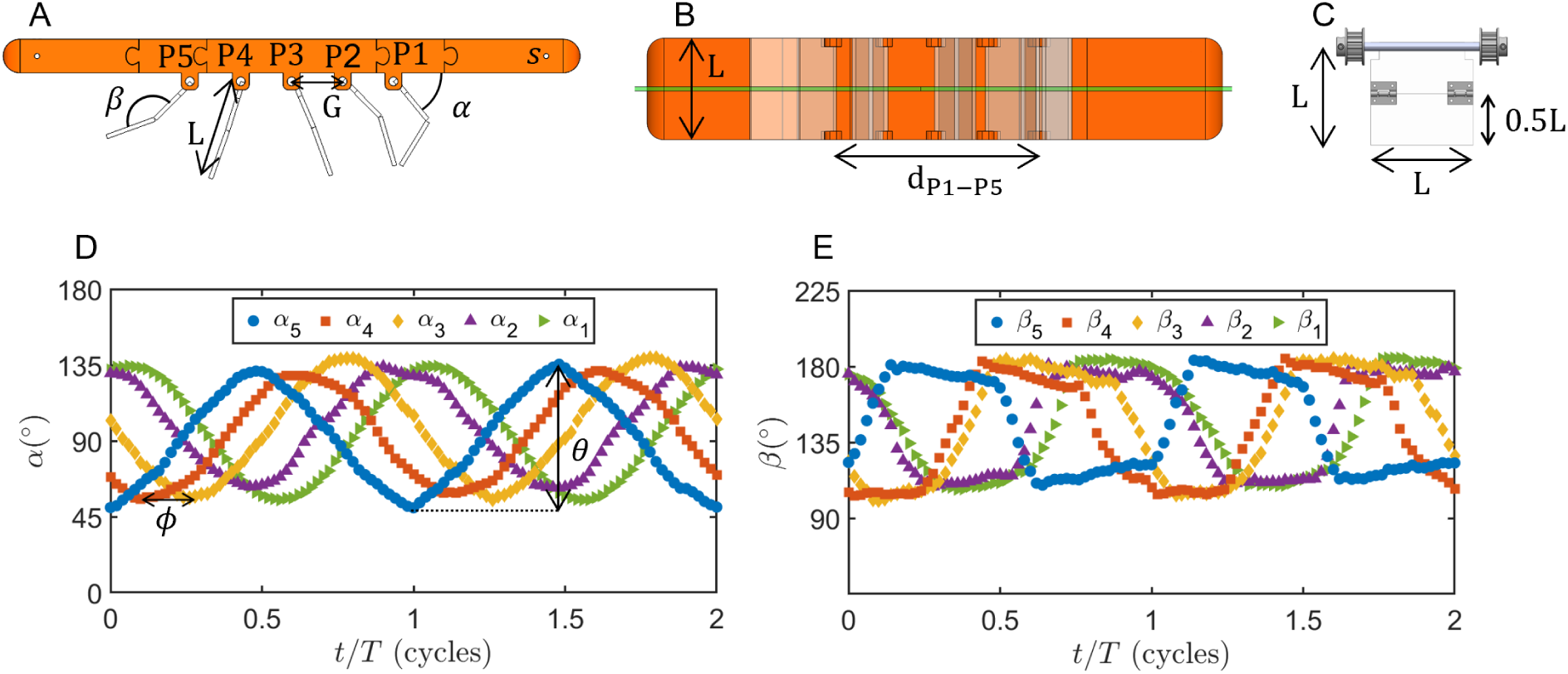
Experimental setup used in this study. A) Robotic model positioned in the aquarium. B) Robotic model with the positions of paddle 1 (P1) and paddle 5 (P5) indicated, as well as paddle length *L*, inter-paddle gap *G* and P1 − P5 distance *d*_P1_*_−_*_P5_. C) Perpendicular view of the paddle. Paddles have an aspect ratio (i.e., length/width) of 1, with a passively rotating hinge joint at the mid-point along the paddle length. D) Tracked paddle angular positions, *α* at *Re*_L_ = 54, 724. E) tracked hinge angles, *β* at *Re*_L_ = 54, 724.

The model was adapted with a generalized flat-plate body to allow for studies of the hydrodynamic scalability of metachronal rowing without confounding factors that result from species-specific kinematics and species-specific morphology (body shapes, appendage sizes and appendage shapes) [28, 29]. The body was fitted with 5 paddles of length *L* = 76.2 mm and spacing between the roots of the paddles (*G*) equal to 0.5*L* (Figure 1A-B). Each paddle had a hinge halfway down its length (*L/*2 = 38.1 mm from the root, Figure 1C). The hinge allowed for passive (i.e., non-actuated) rotation between internal angles (*β*) of approximately 110*^◦^* to 180*^◦^* according to the fluid dynamic forces acting on the paddles. Hinge angles over two stroke cycles were manually digitized from high-speed recordings using the angle tool in ImageJ (version ij153 [33]), an example of tracked hinge kinematics for *Re*_L_ = 54, 724 are shown in Figure 1E. Hinge angles for the other conditions are included in the supplementary material for this study (Figure S2). Paddles were found to fold early in the recovery stroke into a configuration that minimized surface area perpendicular to the direction of motion of the paddle tip, and to unfold early in the power stroke into a configuration that maximizes surface area perpendicular to the paddle tip motion. This is qualitatively similar to the appendage bending kinematics reported in a variety of metachronally rowing crustaceans including Antarctic krill [5].

### 2.2 Test conditions

Although many organisms that use metachronal rowing show variations in stroke amplitudes, stroke angles and phase lags for paddles at different positions along the body [5, 34, 35], these differences are both species- and behavior-dependent. In this study, a constant stroke amplitude (*θ* = *π/*2 rad) was prescribed for each paddle and a constant phase lag (*ϕ* = 0.15; i.e., 15% of the stroke period) was maintained between all adjacent pairs of paddles on the model across all test conditions (Figure 1D). The stroke period is defined as *T* = 1*/f*, so that the product *ϕT* gives the delay between the movement of adjacent paddles in seconds. Time was nondimensionalized against the stroke period, *τ* = *t/T*.

The robot was submerged in a custom glass aquarium, measuring 244 cm (length) × 65 cm (width) × 77 cm (height), which was filled with three different water-glycerin mixtures. Stroke frequencies were varied from *f* = 1.0 Hz to *f* = 3.0 Hz in steps of 0.5 Hz. For a given stroke frequency, the three mixtures had kinematic viscosity (*ν*) values of 1 mm^2^ s^−1^, 100 mm^2^ s^−1^ and 860 mm^2^ s^−1^. This combination of stroke frequencies and viscosities resulted in 15 *Re*_L_ test conditions, ranging from *Re*_L_ = 21 to *Re*_L_ = 54724 (Table 1). *Re*_L_ was defined according to the following equation:

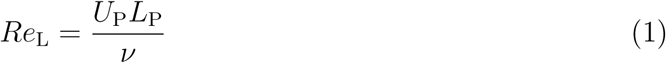

where *U*_p_ is the mean paddle tip speed, which can be calculated as:

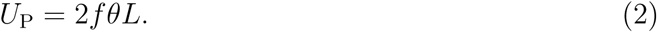

**Table 1.**
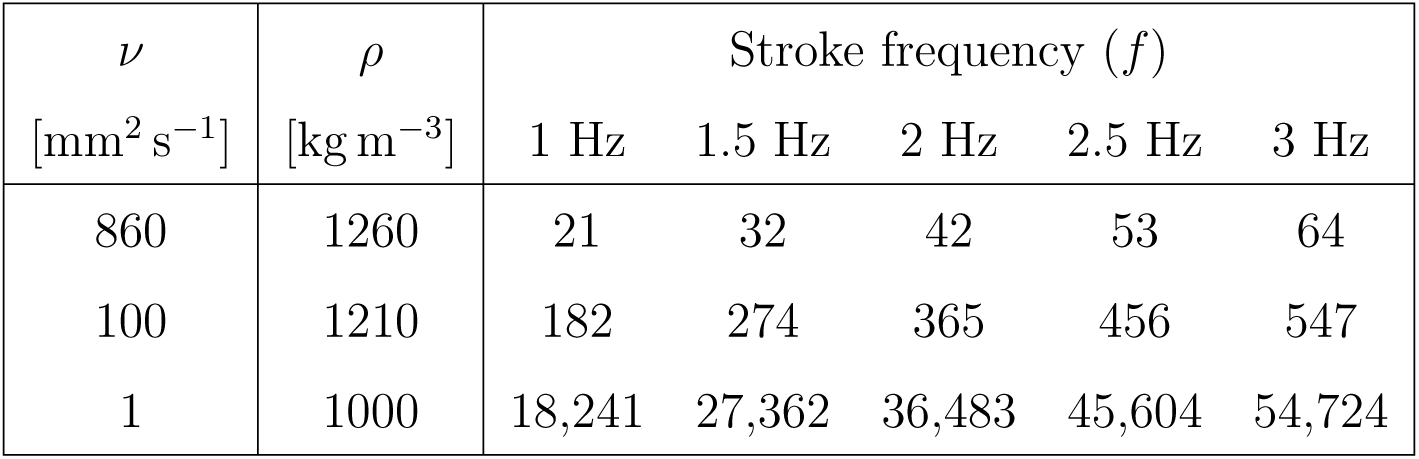
Experimental conditions used to obtain the 15 unique paddle Reynolds numbers (*Re*_L_) examined in this study. Stroke frequency (*f*) and kinematic viscosity of water-glycerin mixture (*ν*) were independently varied to obtain *Re*_L_ ranging from 21 to 54,724.

| $\nu$<br>[mm<sup>2</sup> s<sup>-1</sup>] | $\rho$<br>[kg m<sup>-3</sup>] | Stroke frequency ($f$) | | | | |
| --- | --- | --- | --- | --- | --- | --- |
|  |  | 1 Hz | 1.5 Hz | 2 Hz | 2.5 Hz | 3 Hz |
| 860 | 1260 | 21 | 32 | 42 | 53 | 64 |
| 100 | 1210 | 182 | 274 | 365 | 456 | 547 |
| 1 | 1000 | 18,241 | 27,362 | 36,483 | 45,604 | 54,724 |

Since the swimming animals directly control their appendage kinematics and not their swimming speed, *Re*_L_ is more appropriate than *Re*_B_ for a systematic investigation of the flow physics of metachronal propulsion. However, measurement of *U*_p_ on live organisms can require high-speed, high-resolution video data which is often unavailable from organismal studies. In these studies, *Re*_B_ is often calculated based on the body size and swimming speed according to the following:

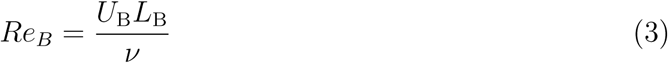

where *L*_B_ is the body length and *U*_B_ is the swimming speed. However, when considering the morphological diversity of the animals that use metachronal rowing, it is apparent that body length and shape will have an effect on the swimming speed that is independent of the hydrodynamic performance of the propulsive movement of the paddling legs.

For the swimming speed experiments, the robot was allowed to swim freely along the length of a 1 m air bearing under its own thrust, while for flow measurements (particle image velocimetry) the robot was tethered in place in order to keep the wake centered in the camera field of view (FOV). During swimming experiments, the air bearing minimized frictional resistance from the mounting system and restricted movement to one dimension.

### 2.3 Swimming speed

Swimming speeds were determined from videos of the self-propelling robotic model (untethered). These videos were recorded using a single highspeed Phantom camera (Miro M110, Vision Research, Wayne, NJ, USA) at a rate of 100 frames per stroke (i.e., 100*f* Hz), where the cameras was positioned such that its sensor plane was parallel to the long-axis of the robot body. The complementary metal oxide semiconductor (CMOS) sensor of the camera was 25.6 × 16.0 mm in size and had a maximum resolution of 1280 × 800 pixels. The camera was fitted with a 60 mm fixed focal-length Nikon Nikkor lens to obtain a FOV measuring 731 × 457 mm^2^ at a spatial resolution of 1.75 pixels/mm. The DLTdv8 tool [36] in Matlab (v9.8; Mathworks Inc, Natick, MA, USA) was used to semi-automatically digitize (with manual intervention to correct tracking errors) the position of one of the mounting screws on the model (labeled “s” in Figure 1A). Numerical differentiation of the digitized position data was performed using a second-order central difference method, and the resulting body speed (*U*_B_) was used for the calculation of the swimming speed based Reynolds number (*Re*_B_, see equation 3). However, since the robotic model has a body geometry of arbitrary length, the appropriate length scale to use for the *Re*_B_ calculation is not obvious. Previous work has been done examining the effect of spacing between the adjacent paddles [28], so we chose the distance between the roots of the first paddle (P1) and the last paddle (P5), *d*_P1_*_−_*_P5_ = 2*L*, to use for the characteristic length since we maintained constant *G/L* = 0.5. This length scale (*d*_P1_*_−_*_P5_) is equal to 4*G* for the 5-paddle model used in this study (see Figure 1B). The robot *Re*_B_ definition is modified from equation 3 in order to account for this change in length scale as:

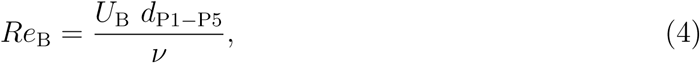

where it is important to note that while the characteristic length (*d*_P1_*_−_*_P5_) is from the model design and *ν* is a prescribed variable (along with *f*, *θ*, *ϕ*), *U*_B_ emerges from the interaction of the paddling-generated flow with the robotic swimming limbs (5 paddles).

### 2.4 Flow visualization

Two-dimensional, two-component, time-resolved particle image velocimetry (PIV) measurements were performed to examine how the flow structures and the wake are modified with varying *Re*_L_. Raw PIV images were recorded using the same high-speed camera as that used for swimming performance measurement, with a 60 mm Nikon Nikkor lens and a spatial resolution of 1.79 pixels/mm, for a total field of view of 715 × 447 mm^2^. DaVis 10 software (DaVis 10.0, LaVision GMBH, Göttingen, Germany) was used to synchronize the camera with a high speed Nd:YLF laser with a wavelength of 527 nm, maximum pulse energy of 30 mJ/pulse and maximum pulse frequency of 10 kHz (Photonics Industries International, Ronkonkoma, NY, USA). The laser beam passed through −10 mm and −20 mm plano-concave cylindrical lenses in series, which spread the beam into a thin (approximately 3 mm) laser sheet. The laser sheet was centered along the width of the model (green line in Figure 1B) and illuminated the full width of the camera FOV. 55 µm diameter polyamide seeding particles (LaVision GMBH, Göttingen, Germany) with specific gravity *SG* = 1.2 were mixed into the three fluids to scatter the laser light for PIV data processing. Average particle image densities *N*_ppp_ were approximately *N*_ppp_ = 0.018 in the water-glycerin mixture, *N*_ppp_ = 0.02 in the pure glycerin, and *N*_ppp_ = 0.026 in water. Cross-correlation analysis of raw PIV image pairs was performed with multiple passes of decreasing interrogation window size, with 1 pass using a window size of 48 × 48 pixels with 50% overlap, and 1 pass using a window size of 24 × 24 pixels with 50% overlap. Vectors were post-processed using a Q-ratio filter to remove vectors with peak Q-ratio < 1.2, a spatial median filter to remove vectors with more than 5 standard deviations difference from neighboring vectors and a smoothing filter over a 3 × 3 grid. Velocity field data containing two-dimensional positions and velocity components were exported for post-processing in Matlab. Raw videos for PIV data analysis at *Re*_L_ 42, 365 and 36483 are provided as electronic supplementary material.

#### 2.4.1 Momentum fluxes

Horizontal and vertical components of the time-averaged momentum flux per unit width of the paddle were calculated paddling wake. Time-varying momentum fluxes were averaged across 15 stroke cycles and nondimensionalized to serve as a surrogate measure of the dimensionless forces (per unit paddle width) imparted by the oscillating paddles to the ambient fluid. The nondimensional cycle-averaged momentum fluxes were defined according to the following equations:

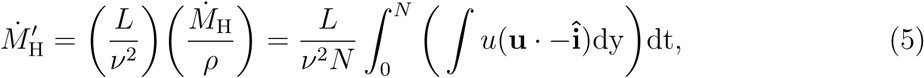

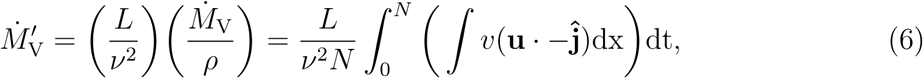

where 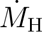 and 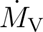 are the horizontal and vertical momentum fluxes, respectively, and *ρ* is the fluid density (values of *ρ* are listed in Table 1). The flow velocity vector is represented by 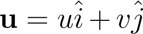, where *u* and *v* are the velocity components along the horizontal and vertical directions, respectively. 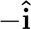 and 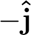 represent the unit normal vectors along the horizontal (directed leftward) and vertical (directed downward), respectively. Time integration is performed over duration *N* = *nT* where *n* = 20 cycles and stroke period *T* is the reciprocal of stroke frequency (*f*). For 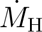, the velocity integral is computed over the entire height of the PIV FOV along a vertical line located at a distance *L* immediately to the left (downstream) of the P5 paddle. For 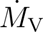, the velocity integral is computed over the entire width of the PIV

FOV along a horizontal line located 2*L* below the roots of the paddles, which is equivalent to distance *L* from the tips of the fully extended paddles (90*^◦^* relative to the horizontal). Diagrams showing the positions at which these calculations were performed are included in the supplementary material, Figure S1.

#### 2.4.2 Dimensionless wake characteristics

Strouhal number (*St*_w_) based on the maximum velocity in the wake (*U*_w_) was also calculated from the cycle-averaged PIV velocity fields. Maximum velocity in the core of the wake jet (*U*_w_) was calculated along the same horizontal line used for the 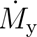 calculation (2*L* vertically down from the paddle roots). The *wake Strouhal number* (*St*_w_) was defined as:

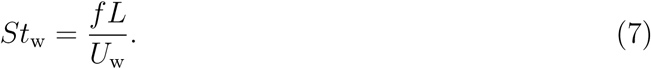

The momentum flux and Strouhal numbers at these locations are useful as measure of hydrodynamic efficiency for swimming. Additionally, the wake also serves an important function to transfer information *in the form of hydrodynamic and chemical signals* between individuals in a swarm or school. In order to quantify how quickly momentum is attenuated in the wake, viscous energy dissipation per unit width (Φ) was calculated in the wake at a region away from the paddles as:

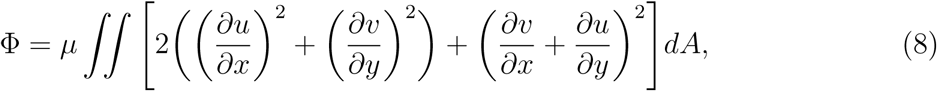

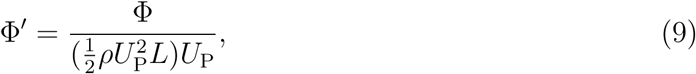

where Φ*^′^* is the nondimensional form of the viscous energy dissipation Φ and 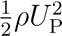 is the dynamic pressure applied on the fluid at the paddle tip due to the paddling motion. Φ*^′^* was calculated over the entire PIV FOV, excluding the region bounded between a distance *L* to the left of P5 to a distance *L* to the right of P1 in the horizontal direction, and from the paddle roots to a distance 2*L* vertically downward from the paddle roots. This region was excluded from the dissipation calculation because within this region, energy is added to the flow by the paddling motion, rather than removed from the flow by viscous dissipation.

#### 2.4.3 Tip vortex tracking

Paddle tip vortices were identified using the swirling strength (*λ_ci_*) criterion [37]. Positive values of *λ_ci_* represent regions of fluid rotation [37], which can be used to identify vortices in a flow field. Vorticity fields were filtered based on *λ*_ci_, and The *circulation* (Γ) of the tip vortex generated on the most posterior/back paddle (P5) during power stroke was calculated. The out-of-plane component of the vorticity vector (*ω_z_*) and *λ_ci_* were calculated in DaVis 10 from the PIV velocity fields. A contour value of *λ_ci_* = 0.0005 was used to define vortices in the PIV velocity fields. Across all test conditions, the maximum value of *λ_ci_* was found to range between 400 and 10,000. Γ represents the strength of the tip vortex, which can also be interpreted as the flux of vorticity in the region occupied by the vortex. The P5 tip vortex was tracked throughout its development and shedding during the power stroke, and its subsequent decay during the recovery stroke.

While vortices were identified automatically using the *λ_ci_* criterion described above, the P5 tip vortex was manually selected from the set of identified vortices. Manual selection was needed to discriminate the P5 tip vortex from the tip vortices generated by the other paddles. Identification of the P5 tip vortex and vorticity integration were performed using a custom Matlab script. Time-varying Γ was calculated as the spatial integral of out-of-plane vorticity within the area occupied by the P5 tip vortex and non-dimensionalized as follows:

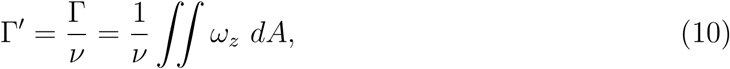

where the nondimensional circulation (Γ*^′^*) is defined as the ratio of instantaneous circulation to the kinematic viscosity of the fluid (*ν*), and *A* is the area of the region occupied by the vortex. As Γ is related to the rotational inertia in a vortex and *ν* is related to the viscous forces in the same flow, the peak value of the Γ*^′^* was defined as the *vortex Reynolds number*, *Re*_Γ_ [38]:

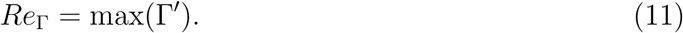

The sensory hairs on some crustaceans are arranged in a way that they can be useful for sensing vorticity in the form of shear or localized rotation in a flow [39]. In this context, it can be helpful to know how the vortex propagation changes with *Re*_L_. The *vortex propagation speed* (*U*_V_) was defined as:

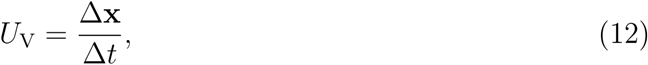

where Δ**x** is the net distance traveled by the vortex between its initial formation and eventual dissipation over the lifetime of the vortex (Δ*t*). The geometric centroid of the vortex is calculated by averaging the positions of each of its discrete elements:

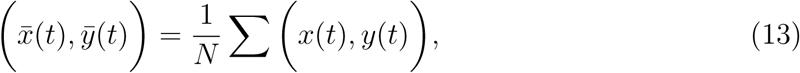

where 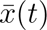 and 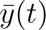 represent the horizontal and vertical positions of the vortex centroid at time *t*, respectively. *N* is the total number of discrete grid points (dependent on vortex size and PIV-based velocity vector spacing) located within the vortex and *x*(*t*) and *y*(*t*) represent the two-dimensional positions of each individual element. The vortex propagation speed can be nondimensionalized against the paddle tip speed to examine vortex propagation efficiency across varying *Re*_L_. This takes a familiar, Strouhal number-like definition, which is termed herein as the *vortex Strouhal number*, *St*_v_:

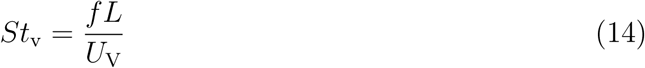

Since *U*_V_ is a term several steps derived from the experimental data, it is more prone to contamination from experimental and numerical noise. However, *U*_V_ provides a way to quantify vortex propagation efficiency that can be of use in studies of the sensory ecology of crustaceans [39].

## 3 Results

### 3.1 Metachronal Paddling Performance

Swimming Reynolds number *Re*_B_ and paddling Reynolds number *Re*_L_ were calculated from previously published data for a variety of morphologically diverse paddling organisms including crustaceans (Subphylum: *Crustacea*), ctenophores (Phylum: *Ctenophora*), annelids (Phylum: *Annelida*) and a single-celled paramecium (Phylum: *Ciliophora*) [5, 7, 15, 20, 34, 35, 40–56]. The two Reynolds numbers were assumed to follow a power law scaling, 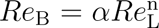, and were found to be approximately linearly correlated (*α* = 3.53, *n* = 1.03). Among the organisms considered, *Re*_L_ is a strong predictor of *Re*_B_ (*R*^2^ = 0.886, *p <* 1×10^−21^, log scale). This scaling is indicated by a dashed line in Figure 3A. The individual points from this dataset are included in a the supplementary material in Table S1. Since the scaling exponent n is approximately 1, the scaling constant *α* is therefore approximately equal to *U*_B_*/U*_P_ and directly proportional to the advance ratio. Advance ratio is a common measurement of hydrodynamic “efficiency” used in swimming studies. However, these quantities (*α* and advance ratio) are poorly suited for direct comparison across species that have widely varying body morphologies and swimming behaviors, and therefore is not considered in this study.

For swimming performance experiments, the robotic model was mounted to a linear air bearing to allow for 1D swimming movement with low frictional resistance. The model was operated with stroke frequencies of 1.5-3 Hz in steps of 0.5 Hz and with phase lags between adjacent paddles of 15% of the stroke period. These kinematics corresponded to Reynolds numbers ranging from *Re*_L_ = 32 to *Re*_L_ = 54724 (Table 1). Data was not recorded for a stroke frequency of 1 Hz, because the model was unable to overcome the initial static friction imposed by the electrical cables (used to provide control inputs and power to the motors) resting on the support frame that enclosed the aquarium. However, for stroke frequencies of *f* = 1.5 Hz and greater, the robot was able to swim under its own power with cycle-averaged swimming speed *U*_B_. Swimming speed-based Reynolds number, *Re*_B_ (equation 3) as a function of *Re*_L_ is shown in Figure 3A (black squares). For the robotic model, the power law scaling 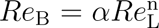 was found to have scaling constant *α* = 0.370 and scaling exponent n = 1.00 (dotted and dashed line in Figure 3). In the controlled robotic experiment, *R*^2^ = 0.995 and *p <* 1 × 10^−12^, which indicates that the linear relationship is due to unchanging physics in the near-wake rather than to selective behavioral differences in the differing organisms. This supports the hypothesis that metachronal paddling is a robust locomotion strategy that can be effectively employed by organisms both small and large, as well as providing support for the application of metachronal paddling as a viable propulsion system for bio-inspired underwater vehicles. Slight deviations from the overall linear trend at the lowest stroke frequencies is again likely due to resistance from the power cables.

In addition to the Reynolds number comparison, nondimensional momentum fluxes in the the horizontal (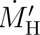) and vertical (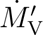) directions were calculated from PIV measurements in the paddling wake. These momentum flux values are directly related to the thrust generation of the metachronal paddling system through the momentum form of the Reynolds transport equation. From equations 5 and 6, it can be seen that 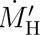 and 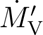 are expected to scale with (*U/ν*)^2^, which is proportional to 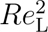. Therefore, if the metachronal paddling system maintains its force generation efficiency across scales in both the thrust (horizontal) and lift (vertical) directions then the relationship 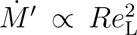 would be expected to hold true. Dimensionless horizontal momentum flux 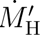 was found to scale with 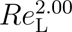, while dimensionless vertical momentum flux 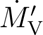 was found to scale with 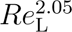. These scaling exponents support that while horizontal force is maintained across scales, vertical force production drops for *Re*_L_ < 42. This indicates that at the smallest scales, metachronal paddling may be less useful for weight support in organisms that are denser than the surrounding fluid environment, although it is still useful for thrust generation.

### 3.2 Hydrodynamics of Metachronal Paddling

Particle image velocimetry was used to measure fluid flow in the wake of the paddling robot. Velocity vector fields overlaid with vorticity contours are shown in Figure 2 for *Re*_L_ = 21 and *Re*_L_ = 54724. Counterclockwise vorticity is indicated in red, while clockwise vorticity is shown in blue. Black contours indicate regions bounded by the vortex identification criterion *λ*_ci_ *>* 0.0005 and identify vortices rotating in the direction according to the sign of the bounded vorticity. The time points are defined according to the angular position of the P5 paddle, which leads the metachronal wave. *τ* = 0 represents the stroke reversal to begin the power stroke, *τ* = 0.25 represents the maximum angular velocity during the power stroke, *τ* = 0.5 represents the stroke reversal to begin the recovery stroke and *τ* = 0.75 the maximum angular velocity of the paddle during the recovery stroke.

**Figure 2.**
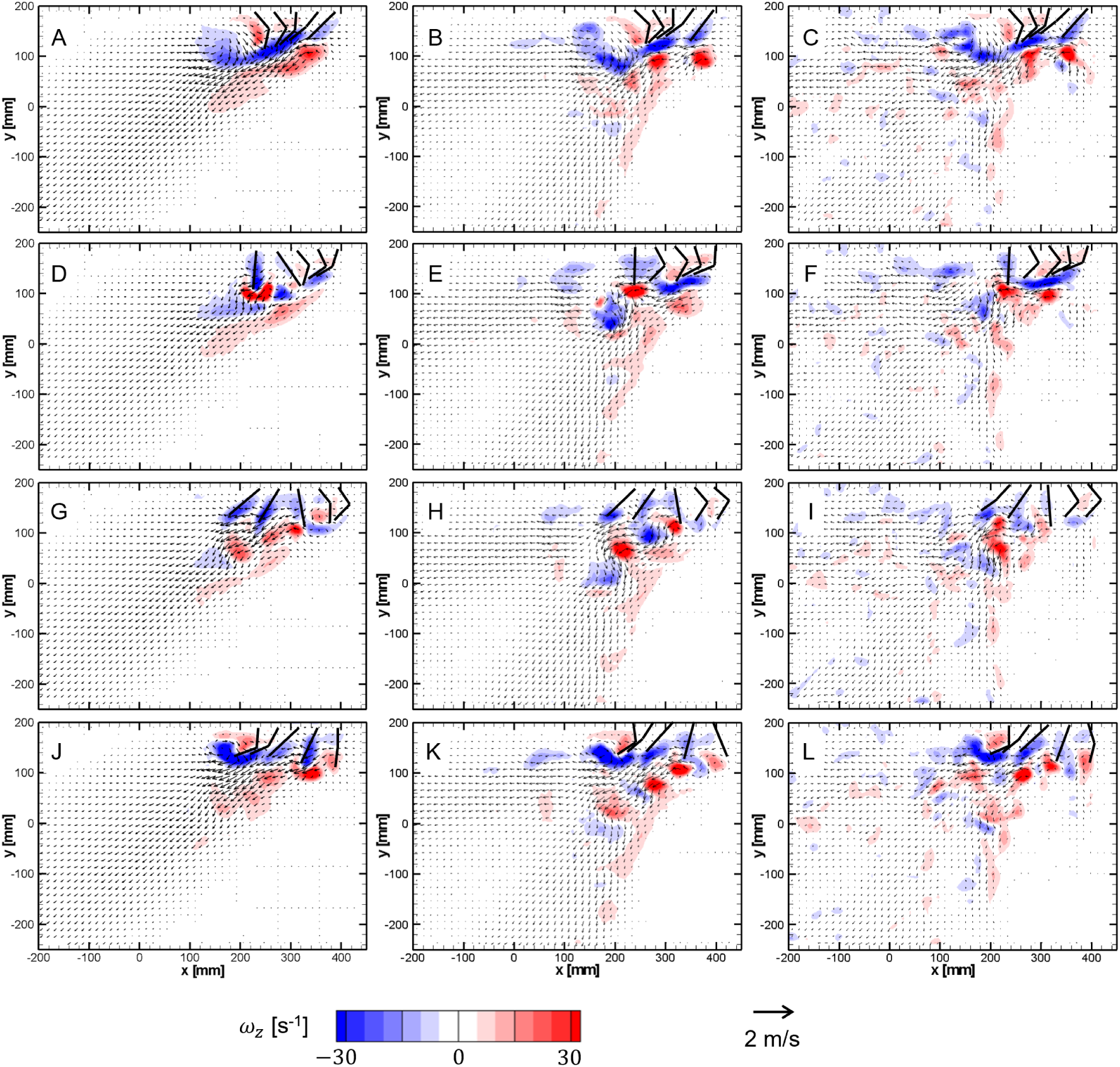
Flow field characteristics. Comparison of the velocity fields (vector arrows) and vorticity contours (colors) at *Re*_L_ = 42, *Re*_L_ = 365 and *Re*_L_ = 36, 483.

**Figure 3.**
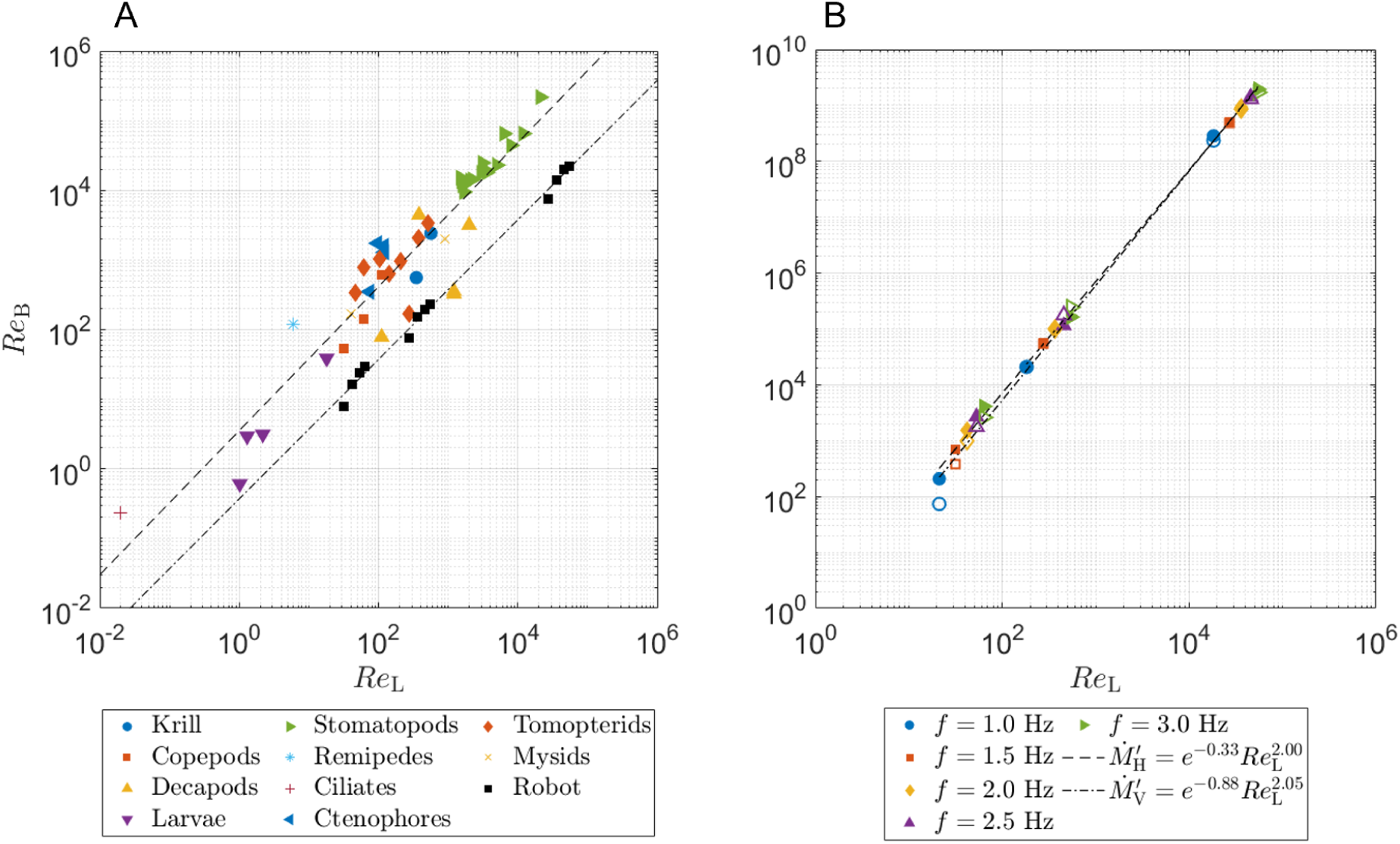
Metachronal paddling performance. A) Comparison of the swimming and paddle Reynolds numbers. The dashed line represents the best fit for the organismal data (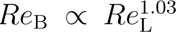). The dashed and dotted line represents the best fit for the robot data (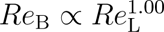). B) Nondimensional momentum flux as a function of paddle Reynolds number in the horizontal (solid markers, dashed line, 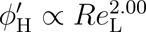) and vertical (hollow markers, dashed and dotted line, 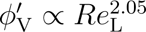) directions.

As time progresses through the stroke cycle, during each paddle’s power stroke (which is defined by a clockwise rotation of the paddles from the reference point of the high speed camera), a tip vortex with a counterclockwise rotation is formed on the paddle. Likewise during the recovery stroke, a tip vortex is formed on the P5 paddle with a clockwise direction of rotation. However, the metachronal motion prevents the sustentation of large tip vortices on the individual forward paddles. It can be seen from Figure 2 that the small-scale wakes generated by the individual paddles interact to form a large-scale wake. This large-scale wake has the characteristic of a periodically oscillating jet, the strength and direction of which depend on the Reynolds number. The tip vortices that form during the recovery strokes of adjacent paddles tend to merge into a single shear layer, contributing to this large-scale wake. This is likely due to the bending motion of the hinges on the paddles, because in contrast to the tip vortices generated during the recovery stroke, it is generally possible to distinguish between the power stroke tip vortices (the vortices that rotate counterclockwise) throughout their entire history, including their generation, shedding and decay.

#### 3.2.1 Tip vortex dynamics

The wake near to the paddles is dominated by vortices that form on the paddle tips during the power and recovery strokes. These paddle-tip vortices can be seen on the tip of the P5 paddle during the power stroke in Figure 2 (parts D, E and F). Circulation was calculated as the integral of the vorticity inside the *λ*_ci_ < 0.001 contour that forms behind the P5 paddle during power stroke. The circulation data was averaged across 5 paddling cycles, and dimensional circulation for *Re*_L_ ∼ *O*(10^1^) and *Re*_L_ ∼ *O*(10^4^) is shown in Figure 4A and Figure 4B, respectively. Circulation is seen to increase with increasing paddle stroke frequency *f*.

**Figure 4.**
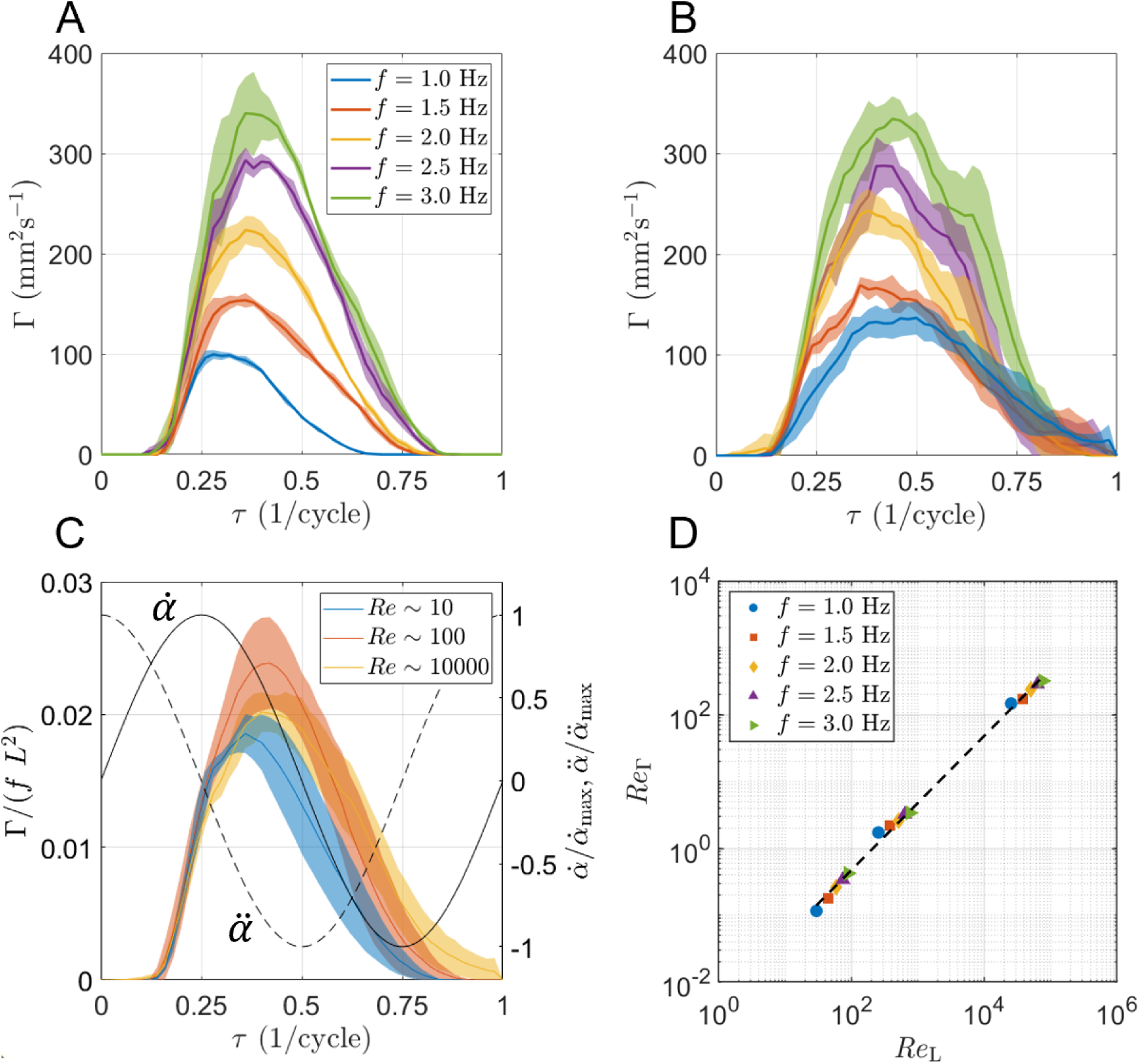
Tip-vortex circulation. A) Circulation in the power stroke tip vortex over time for each frequency at *ν* = 860 mm s^−2^. Lines indicate the mean value across 5 trials. Patches indicate the standard deviation. B) Circulation in the power stroke tip vortex with *ν* = 1 mm s^−2^. C) Tip vortex circulation nondimensionalized to remove the effect of varying kinematics. The vortex development is similar across all conditions, but the decay and peak circulation change slightly with Reynolds number. D) *Re*_Γ_ as a function of *Re*_L_. *Re*_Γ_ scales linearly with *Re*_L_.

The tip vortex circulation can be nondimensionalized for comparison across different conditions by removing either the effect of paddle kinematics (Figure 4C), or the effect of viscosity (Figure 4A). From the scaling in Figure 4C, the tip vortex circulation largely collapses to similar values across differing frequencies and viscosities, although increasing *Re*_L_ across orders of magnitude generally results in a slightly delayed shedding of the tip vortex (peak shifts to the right). In general, this nondimensional tip vortex circulation follows the kinematics of the P5 paddle during the power stroke with a slight delay. The paddles reach a maximum angular acceleration during the power stroke at *τ* = 0.125, a maximum angular velocity at *τ* = 0.25, and a maximum angular deceleration at *τ* = 0.375. The tip vortex begins to strengthen rapidly at *τ* ≈ 0.14 across all *Re*_L_, while there is a slight delay in the vortex shedding with increasing *Re*_L_, with Γ*/*(*f L*^2^) reaching a maximum at *τ* ≈ 0.36 for *Re*_L_ ∼ *O*(10^1^), *τ* ≈ 0.40 for *Re*_L_ ∼ *O*(10^2^), and *τ* ≈ 0.42 for *Re*_L_ ∼ *O*(10^4^). After the vortex is shed from the paddle tip, the circulation begins to decay due to viscous effects in the far wake. The shed vortex decays slower at higher *Re*_L_, but after being shed the vortex no longer contributes directly to thrust generation on the paddle and is not considered here.

Dimensionless circulation was also calculated with viscous forces removed (Γ*^′^*), and the peak value of Γ*^′^* was found to increase both with increasing stroke frequency and with decreasing viscosity. The vortex Reynolds number *Re*_Γ_ was defined to be the peak value of Γ*^′^* for each test condition. Figure 4D shows *Re*_Γ_ as a function of *Re*_L_. The scaling was found to follow a power law 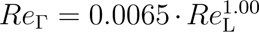, *R*^2^ = 0.999. This 1:1 power scaling of *Re*_Γ_ to *Re*_L_ further supports the idea that metachronal paddling maintains propulsive efficiency across the tested range of *Re*_L_ since circulation in an attached vortex is directly related to thrust generation.

#### 3.2.2 Strouhal numbers and viscous dissipation

Although the near-wake dynamics are most important for the thrust generation, the far-wake dynamics are important for sensing hydrodynamic signals for swarming behaviors and predation avoidance. Dimensionless viscous energy dissipation Φ*^′^* and Strouhal numbers based on the maximum velocity in the core of the cycle-averaged wake jet (*St*_w_) and on the advection velocity of the P5 paddle tip vortex after vortex shedding (*St*_v_) were calculated. As *Re*_L_ increases, so does the ratio of inertial to viscous forces, resulting in a decrease in viscous dissipation. Φ*^′^* was found to follow a scaling law of 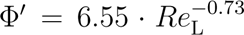, with Φ*^′^* increasing with decreasing *Re*_L_ until *Re*_L_ = 42 (Figure 5). This indicates that viscous effects attenuate the hydrodynamic signals in the wake much more quickly at low *Re*_L_ than at higher *Re*_L_.

**Figure 5.**
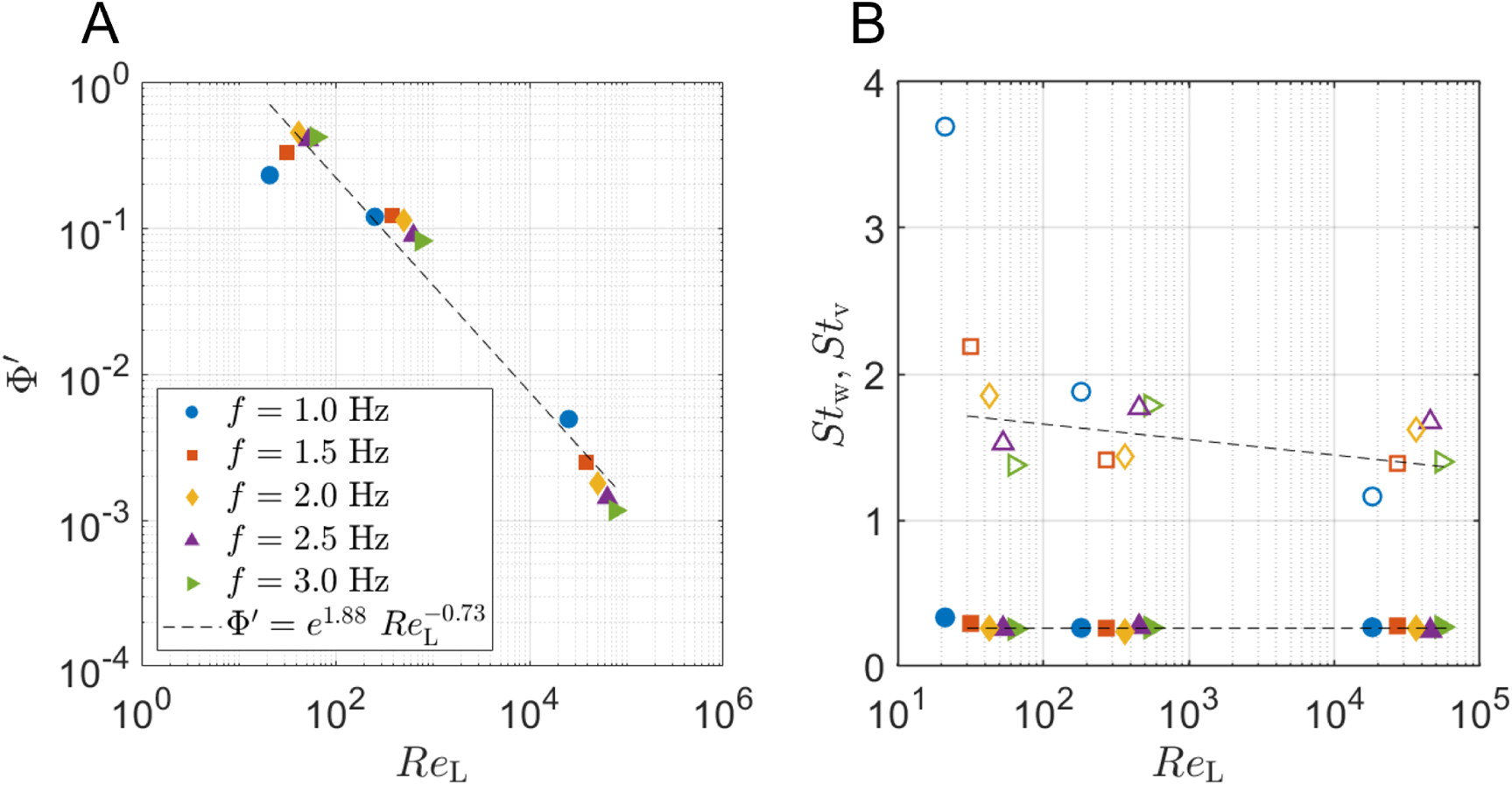
Attenuation of hydrodynamic signals in the metachronal paddling wake. A) Nondimensional viscous energy dissipation indicates that total kinetic energy in the wake is reduced at low *Re*_L_. B) Strouhal numbers based on the velocity in the core of the wake jet (*St*_w_, solid markers) and on vortex transport (*St*_v_, hollow markers) suggest that the core of the wake jet can carry velocity information farther than the vorticity in paddle tip vortices after shedding.

The wake Strouhal number, *St*_w_ was found to be nearly constant at *St*_w_ = 0.26 in almost the entire range of *Re*_L_ tested, 42 <= *Re*_L_ <= 54724. This is shown in Figure 5. In contrast, the vortex Strouhal number *St*_v_ was found have a slight negative dependence on *Re*_L_, for *Re*_L_ ≥ 42, but to increase rapidly with decreasing *Re*_L_ for *Re*_L_ < 42. Since the Strouhal number can be a measure of how efficiently momentum is transferred from the paddle to the fluid in the wake, a constant or nearly constant value of *St*_w_ and *St*_v_ further indicate that the metachronal paddling system is equally well-suited for generating propulsive momentum for Reynolds numbers across the tested range of 42 ≤ *Re*_L_ ≤ 54724.

## 4 Discussion

In nature, metachronal paddling is used as a propulsion strategy by organisms across a wide range of Reynolds number regimes, ranging from microscopic ciliates that swim in a viscosity-dominated regime at appendage Reynolds number *Re*_L_ ∼ 10^−2^ [45], to large benthic crustaceans in a firmly inertial flow regime with *Re*_L_ ∼ 10^4^ [15, 34]. In this study, we examined how changing the paddle Reynolds number affects swimming speed of the model and momentum flux in surrounding fluid, tip vortex development, and hydrodynamic signals in the far-wake.

To the best of our knowledge, this is the first study considering metachronal paddling across multiple orders of magnitude of *Re*_L_ on a scale of *Re*_L_ ≥ 10^1^. By maintaining the same physical model and prescribed paddle kinematics across these scales, we are able to generalize observations about the metachronal paddling system independent of the morphological and kinematic differences that are present when comparing animal species, in order to determine the physical mechanisms of metachronal paddling across scales.

### 4.1 Swimming performance is independent of Reynolds number

Swimming performance was measured using nondimensional swimming speed (*Re*_B_) and nondimensional momentum flux (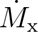 and 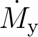). Swimming speed in the metachronally paddling organisms was found to scale as 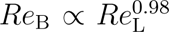, while swimming speed in the robotic model was found to scale as 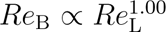. Nondimensional horizontal momentum flux was found to scale as 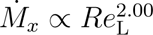. While *Re*_L_ is controlled based on the stroke frequency of the paddle and the viscosity of the fluid, *Re*_B_ and 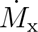 are both emergent from the interactions of the paddles with the fluid. Together, these results indicate that the metachronal paddling strategy is equally effective at converting paddle tip motion into forward swimming motion across all *Re*_L_ scales tested. In contrast, nondimensional vertical momentum flux was found to scale as 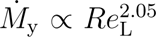. Crustaceans that are denser than water have to generate vertical force to support their weight while swimming in addition to the thrust needed to propel their bodies forward. At high values of *Re*_L_, we find that metachronal paddling can be particularly useful for weight support, with 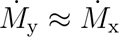 for *Re*_L_ ≥ 182. However, as *Re*_L_ decreases in the range *Re*_L_ ∼ *O*(10^1^), 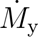 becomes much lower than 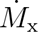, indicating that at low *Re*_L_ the metachronal paddling strategy is not as useful for generating a vertical, weight-supporting force.

### 4.2 Tip vortex development depends primarily on kinematics

In the wake directly behind a paddle performing its power stroke, a paddle-tip vortex forms [14] which results from a low-pressure region on the leeward side of the paddle [13]. In this study, we find that for all *Re*_L_, the circulation in the developing tip vortex closely follows the kinematics of the paddle (Figure 4). The shedding time and peak circulation of the vortex are slightly affected by *Re*_L_, but the initial slope of the increasing portion of the nondimensional circulation versus time graph is unaffected by changing *Re*_L_. This indicates that in the immediate vicinity of the oscillating paddle, the flow is dominated by the paddle rotation, rather than by fluid inertia or viscous damping of the flow. In lift-based propulsive flows, tip vortex circulation is taken to be directly proportional to the lifting force, although this interaction is of a lesser focus in reciprocal or drag-based propulsion. In this study, we show that that the paddle kinematics Reynolds number *Re*_L_ is directly proportional to both the swimming Reynolds number *Re*_B_ and to the tip-vortex circulation based Reynolds number *Re*_Γ_. This indicates that the angular momentum of the paddles and the fluid should be excellent predictors for propulsive thrust and swimming speed, which could allow for the future development of a predictive model of metachronal paddling thrust and swimming performance based on kinematics and tip vortex circulation.

### 4.3 Viscous dissipation attenuates signals in the far wake

Although the flow in the immediate vicinity of the paddles is found to depend directly on the paddle kinematics, the far-wake, is found to attenuate quickly at *Re*_L_ < 42. This finding is significant because information carried in hydrodynamic and chemical signals in the far-wake are used for communication and sensing in many crustaceans. Male copepods are able to follow chemical trails of potential mates during mate-tracking behaviors, but use hydrodynamic signals to detect when they are near to the potential mate [4]. Pacific krill (*Re*_L_ ≈ 350, [20]) commonly form densely packed, unorganized swarms, but have not been observed to form organized schools, while Antarctic krill (*Re*_L_ ≈ 566, [5]) are able to form swarms as well as highly organized, fast-moving schools. This has been proposed as a result of the relatively longer extent of the paddling wake of the larger *E. superba*, which is supported by the findings of this study. Viscous energy dissipation in the far-field of the metachronal paddling wake was found to scale with *Re*^0.73^, which indicates that at higher *Re*_L_ the hydrodynamic signals in the wake are less attenuated and can be more easily sensed by conspecific individuals.

Additionally, we find that the wake-based Strouhal number in the paddling wake is independent of *Re*_L_. For most swimming animals, Strouhal number is defined based on the ratio of propulsive appendage tip speed (fins, flukes, etc.) to swimming speed. Strouhal number is often used as a measurement of how efficiently momentum is transferred from the propulsive appendages (paddles in this study, but often legs, fins, or flukes in animals) to the paddling wake. For streamlined, forward-swimming animals with densities similar to the surrounding fluid, it can be appropriate to consider the swimming speed of the animal for the calculation of the Strouhal number. However, this definition is only applicable to particular swimming gaits and behaviors. For example, for hovering behaviors Strouhal number based on swimming speed is undefined as the swimming speed is placed in the denominator in the Strouhal number calculation. Therefore, using the velocity in the core of the time-averaged wake jet, we find that the value of *St*_w_ = 0.26, which is well within the range of 0.2 <= *St <*= 0.4 that has been reported for a wide variety of animals in cruising behaviors [57], as well as the range of *St* seen emerging from classic physics, such as flow past a cylinder [58]. This further supports that the metachronal paddling mechanism is robust across several orders of magnitude of Reynolds numbers.

### 4.4 Limitations

In order to examine the effects of scale on the metachronal paddling propulsion system, we adopted a generalized robotic platform that neglect much of the species-specific morphological and kinematic variation on the paddling propulsion strategy seen in nature. The legs of metachronal swimming animals come in a variety of different shapes, ranging from nearly solid, overlapping plates such as in mantis shrimp [34] to long, narrow structures fringed with long setae as in amphipods [59]. Additionally, these legs can be segmented, flexible, or stiff. In order to make direct comparisons between the different conditions in this study, we consider only solid, rectangular paddles with an aspect ratio of 1 (paddle width / paddle length) that are segmented halfway down the length. The hinge at the paddle segment allows passive rotation between angles of 100 deg ≤ *β* ≤ 180 deg. Previous studies have found weak flexor muscles that can help the swimming legs in some crustaceans to bend during the recovery stroke, but the unbending of the leg during the power stroke is believed to be dominated by the fluid resistance [59]. Additionally, the paddle stroke kinematics used by the animals vary depending on species and behaviors, with differing stroke orientations and stroke angles for each leg, differing numbers of legs and even time-varying phase lag between paddles such as in the hybrid metachrony used by escaping copepods and mantis shrimp [44, 60]. The paddle kinematics of the robotic model are generalized to a simple harmonic motion, with each paddle prescribed identical but phase-shifted kinematics.

## 5 Conclusions

The propulsive performance of the metachronal paddling system is found to be robust to changes in paddling Reynolds number (*Re*_L_) across several orders of magnitude from *Re*_L_ ∼ 10 to *Re*_L_ ∼ 10, 000. Swimming Reynolds number (*Re*_B_) and tip vortex Reynolds number (*Re*_Γ_) were found to scale linearly with *Re*_L_, while horizontal momentum flux (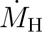) scales quadratically with *Re*_L_ and wake Strouhal number (*St*_w_) remains nearly constant (*St*_w_ = 0.26) over the range of *Re*_L_ from 42 to over 54724. These results indicate that the metachronal paddling strategy is equally effective at producing propulsive forces at high *Re*_L_ as it is at low *Re*_L_.

Both the Reynolds number based on body length and swimming speed (*Re*_B_), and the Reynolds number based on circulation in from the tip vortex of the last paddle (*Re*_Γ_) were found to scale linearly with the Reynolds number based on paddle tip-speed (*Re*_L_), indicating that size and stroke frequency are limited by the constraints of the paddling system, rather than by fluid forces. This is further supported by the collapse of the ascending portion of the Γ*/*(*f L*^2^) curve in time along a single line for all conditions of *Re*_L_. Together, all these results indicate that the system is robust enough for implementation as a bio-inspired propulsion mechanism in autonomous underwater vehicles much larger than the size of a small marine crustacean, and is likely not a primary limiting factor in the size of crustaceans that use metachronal swimming.

## Supporting information

Supplementary Material Description

Example video used for PIV processing (Re_L = 42)

Example video used for PIV processing (Re_L = 365)

Example video used for PIV processing (Re_L = 36,483)

## 6 Supplementary Information

Example videos used for PIV processing, Reynolds number data associated with organisms included in Figure 3A, diagram showing positions used for flow characterization from PIV flow fields, and tracked bending angles are provided as electronic supplementary material.

## Author Contribution Statement

**MPF:** Conceptualization; Investigation; Methodology; Formal Analysis; Visualization; Writing – original draft; Writing – Review & Editing. **AS:** Conceptualization; Formal Analysis; Funding Acquisition; Methodology; Project Administration; Supervision; Writing – Review & Editing.

## Acknowledgements

None.

## SUPPLEMENTARY MATERIAL

### SUPPLEMENTARY TABLE

**Table S1.**
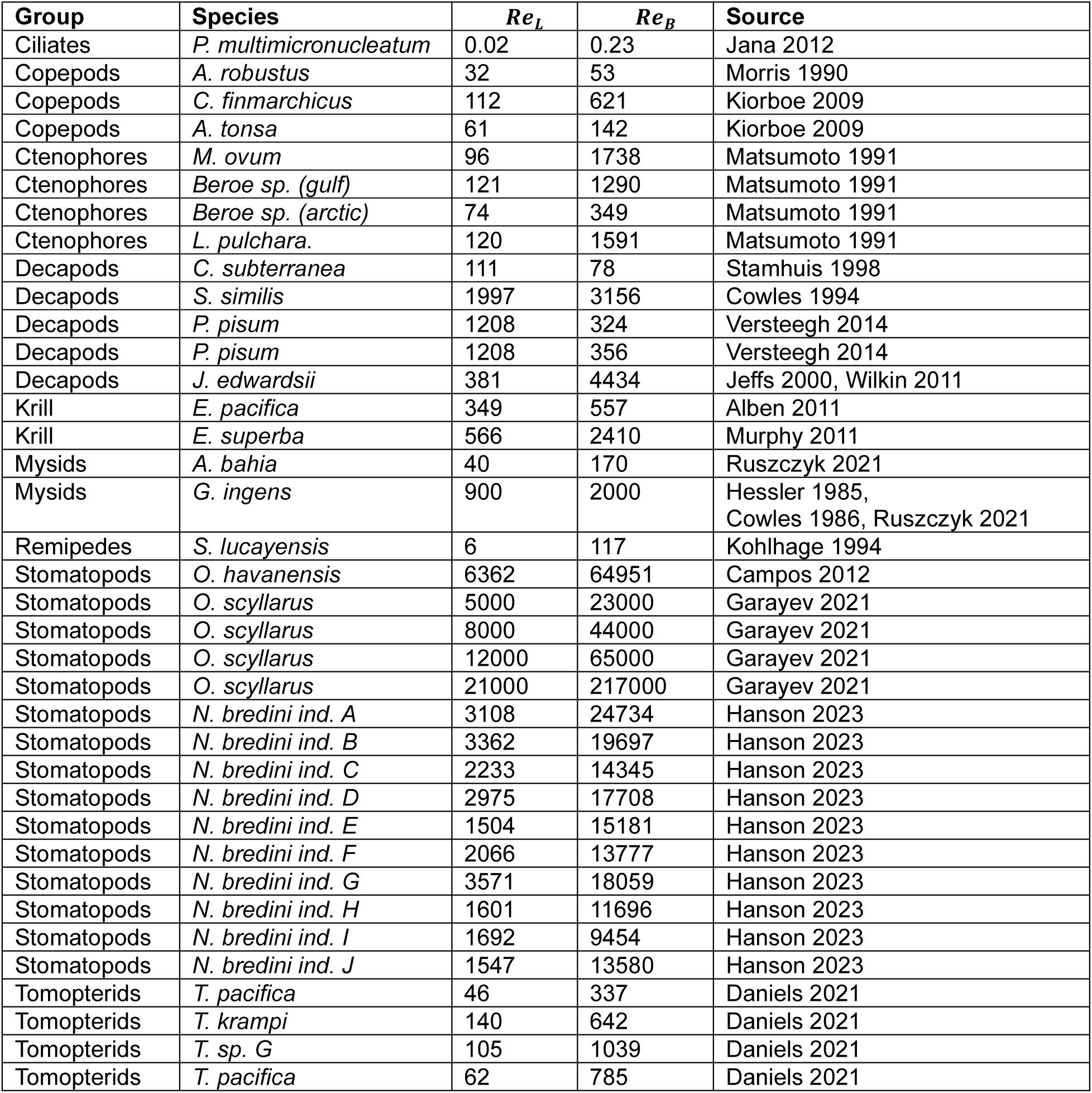
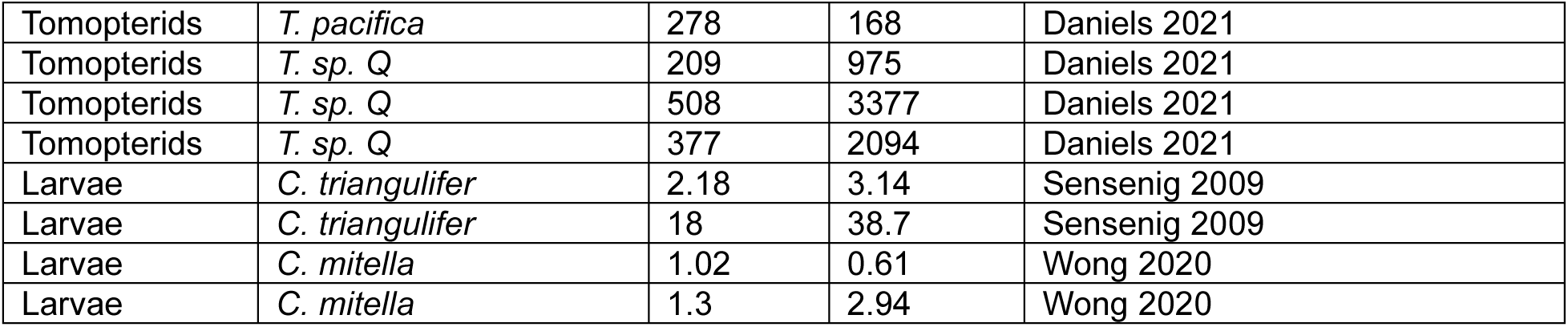
Organismal data included in Figure 3A.

### SUPPLEMENTARY FIGURES

**Figure S1.**
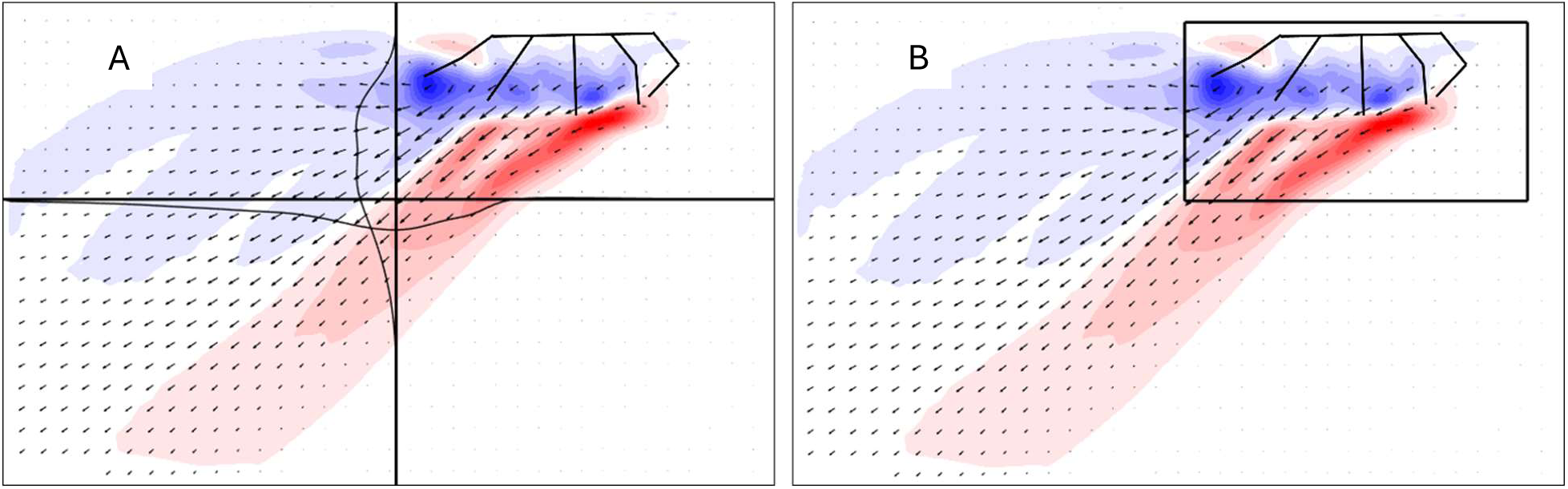
Cycle-averaged PIV data for *Re*_L_ = 64. Positions of the paddles at 50% power stroke are indicated. A) Vertical and horizontal lines where velocity was measured for the 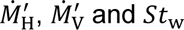 calculations are indicated by bold black lines. The narrower lines indicate the velocity profile normal to the lines. B) Box indicating the region excluded from the Φ, calculation.

**Figure S2.**
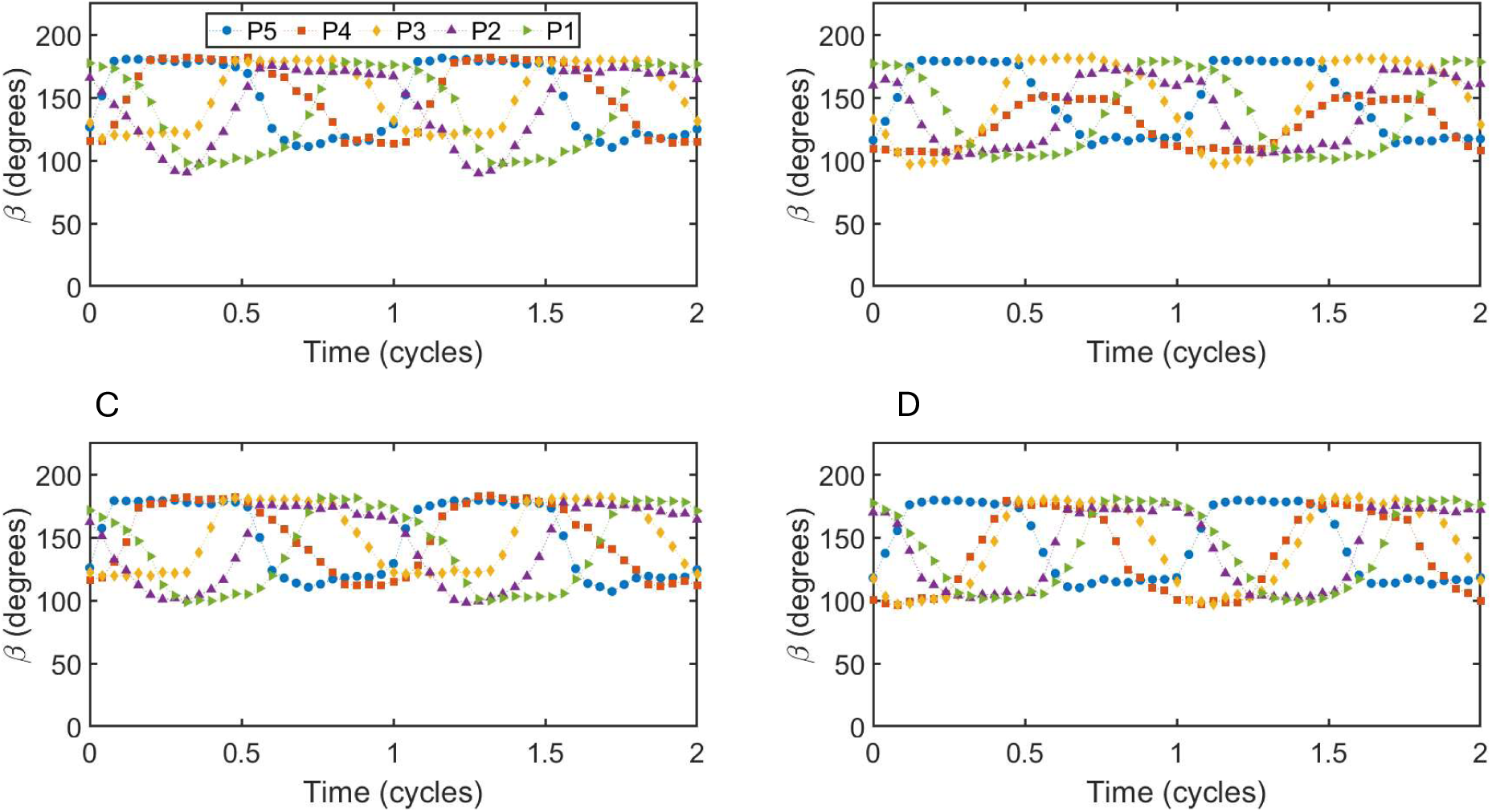
Tracked paddle bending angles for A) *Re*_L_ = 32, B) *Re*_L_ = 27,362, C) *Re*_L_ = 64 and D) *Re*_L_ = 54,724.

### SUPPLEMENTARY VIDEOS

**Video 1.** Example video used for PIV processing for the case of *Re*_L_ = 42. The video has been converted from cine to mp4 format and trimmed to show two stroke cycles. The first frame of the video represents the start of power stroke for the P5 paddle.

**Video 2.** Example video used for PIV processing for the case of *Re*_L_ = 365. The video has been converted from cine to mp4 format and trimmed to show two stroke cycles. The first frame of the video represents the start of power stroke for the P5 paddle.

**Video 3.** Example video used for PIV processing for the case of *Re*_L_ = 36,483. The video has been converted from cine to mp4 format and trimmed to show two stroke cycles. The first frame of the video represents the start of power stroke for the P5 paddle.

### SUPPLEMENTARY DATA

**SupplementaryData.zip:** compressed (ZIP) folder containing the following comma separated values (CSV) files:

**ReL32_Betas.csv:** Tracked paddle bending angle (*β*) versus dimensionless time data for *Re*_L_ = 32.

**ReL64_Betas.csv:** Tracked paddle bending angle (*β*) versus dimensionless time data for *Re*_L_ = 64.

**ReL27362_Betas.csv:** Tracked paddle bending angle (*β*) versus dimensionless time data for *Re*_L_ = 27,362.

**ReL54724_Betas.csv:** Tracked paddle bending angle (*β*) versus dimensionless time data for *Re*_L_ = 54,724.

