## Supplementary Material Description for "Metachronal rowing provides robust propulsive performance across four orders of magnitude variation in Reynolds number"

---

### SUPPLEMENTARY TABLE

**Table S1.** Organismal data included in Figure 3A.

| Group | Species | $Re_L$ | $Re_B$ | Source |
|---|---|---|---|---|
| Ciliates | *P. multimicronucleatum* | 0.02 | 0.23 | Jana 2012 |
| Copepods | *A. robustus* | 32 | 53 | Morris 1990 |
| Copepods | *C. finmarchicus* | 112 | 621 | Kiorboe 2009 |
| Copepods | *A. tonsa* | 61 | 142 | Kiorboe 2009 |
| Ctenophores | *M. ovum* | 96 | 1738 | Matsumoto 1991 |
| Ctenophores | *Beroe sp. (gulf)* | 121 | 1290 | Matsumoto 1991 |
| Ctenophores | *Beroe sp. (arctic)* | 74 | 349 | Matsumoto 1991 |
| Ctenophores | *L. pulchara.* | 120 | 1591 | Matsumoto 1991 |
| Decapods | *C. subterranea* | 111 | 78 | Stamhuis 1998 |
| Decapods | *S. similis* | 1997 | 3156 | Cowles 1994 |
| Decapods | *P. pisum* | 1208 | 324 | Versteegh 2014 |
| Decapods | *P. pisum* | 1208 | 356 | Versteegh 2014 |
| Decapods | *J. edwardsii* | 381 | 4434 | Jeffs 2000, Wilkin 2011 |
| Krill | *E. pacifica* | 349 | 557 | Alben 2011 |
| Krill | *E. superba* | 566 | 2410 | Murphy 2011 |
| Mysids | *A. bahia* | 40 | 170 | Ruszczyk 2021 |
| Mysids | *G. ingens* | 900 | 2000 | Hessler 1985, Cowles 1986, Ruszczyk 2021 |
| Remipedes | *S. lucayensis* | 6 | 117 | Kohlhage 1994 |
| Stomatopods | *O. havanensis* | 6362 | 64951 | Campos 2012 |
| Stomatopods | *O. scyllarus* | 5000 | 23000 | Garayev 2021 |
| Stomatopods | *O. scyllarus* | 8000 | 44000 | Garayev 2021 |
| Stomatopods | *O. scyllarus* | 12000 | 65000 | Garayev 2021 |
| Stomatopods | *O. scyllarus* | 21000 | 217000 | Garayev 2021 |
| Stomatopods | *N. bredini ind. A* | 3108 | 24734 | Hanson 2023 |
| Stomatopods | *N. bredini ind. B* | 3362 | 19697 | Hanson 2023 |
| Stomatopods | *N. bredini ind. C* | 2233 | 14345 | Hanson 2023 |
| Stomatopods | *N. bredini ind. D* | 2975 | 17708 | Hanson 2023 |
| Stomatopods | *N. bredini ind. E* | 1504 | 15181 | Hanson 2023 |
| Stomatopods | *N. bredini ind. F* | 2066 | 13777 | Hanson 2023 |
| Stomatopods | *N. bredini ind. G* | 3571 | 18059 | Hanson 2023 |
| Stomatopods | *N. bredini ind. H* | 1601 | 11696 | Hanson 2023 |
| Stomatopods | *N. bredini ind. I* | 1692 | 9454 | Hanson 2023 |
| Stomatopods | *N. bredini ind. J* | 1547 | 13580 | Hanson 2023 |
| Tomopterids | *T. pacifica* | 46 | 337 | Daniels 2021 |
| Tomopterids | *T. krampi* | 140 | 642 | Daniels 2021 |
| Tomopterids | *T. sp. G* | 105 | 1039 | Daniels 2021 |
| Tomopterids | *T. pacifica* | 62 | 785 | Daniels 2021 |

| | | | | |
|---|---|---|---|---|
| Tomopterids | *T. pacifica* | 278 | 168 | Daniels 2021 |
| Tomopterids | *T. sp. Q* | 209 | 975 | Daniels 2021 |
| Tomopterids | *T. sp. Q* | 508 | 3377 | Daniels 2021 |
| Tomopterids | *T. sp. Q* | 377 | 2094 | Daniels 2021 |
| Larvae | *C. triangulifer* | 2.18 | 3.14 | Sensenig 2009 |
| Larvae | *C. triangulifer* | 18 | 38.7 | Sensenig 2009 |
| Larvae | *C. mitella* | 1.02 | 0.61 | Wong 2020 |
| Larvae | *C. mitella* | 1.3 | 2.94 | Wong 2020 |

### SUPPLEMENTARY FIGURES

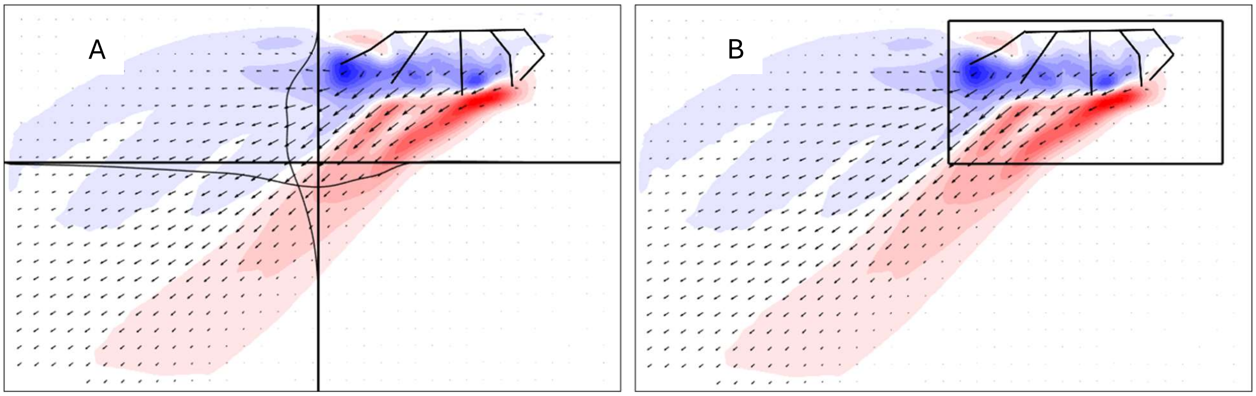


**Figure S1.** Cycle-averaged PIV data for $Re_{\mathrm{L}} = 64$. Positions of the paddles at 50% power stroke are indicated. A) Vertical and horizontal lines where velocity was measured for the $\dot{M}'_{\mathrm{H}}$, $\dot{M}'_{\mathrm{V}}$ and $St_{\mathrm{w}}$ calculations are indicated by bold black lines. The narrower lines indicate the velocity profile normal to the lines. B) Box indicating the region excluded from the $\Phi'$ calculation.

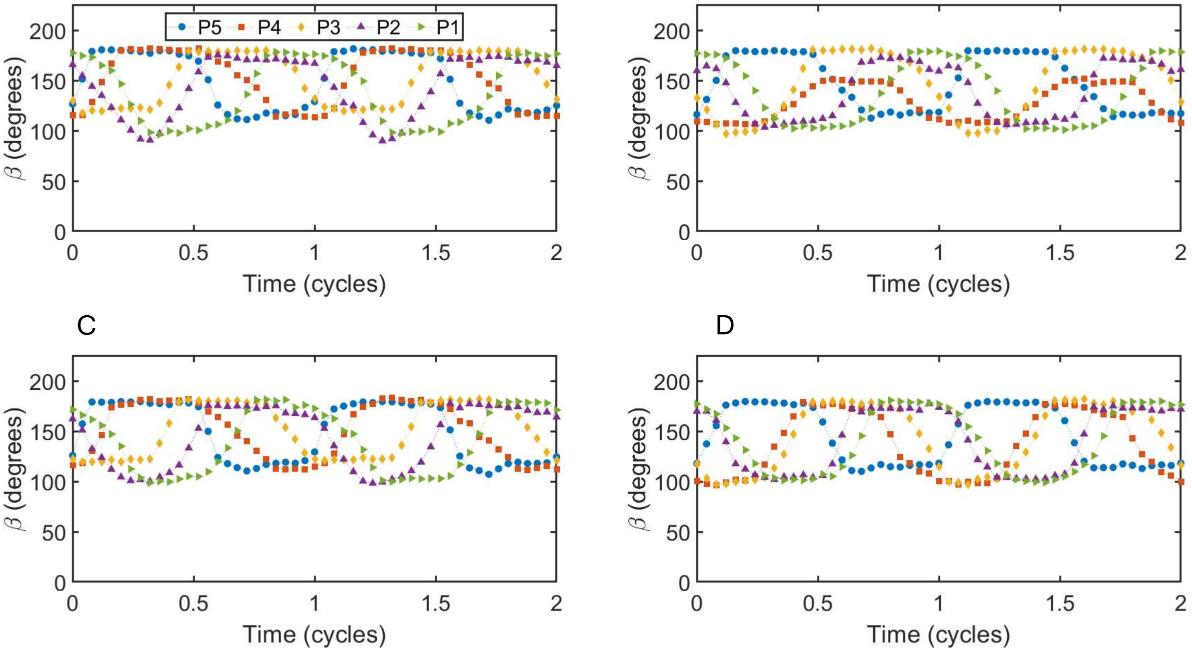


**Figure S2.** Tracked paddle bending angles for A) $Re_{\mathrm{L}} = 32$, B) $Re_{\mathrm{L}} = 27{,}362$, C) $Re_{\mathrm{L}} = 64$ and D) $Re_{\mathrm{L}} = 54{,}724$.

### SUPPLEMENTARY VIDEOS

**Video 1.** Example video used for PIV processing for the case of $Re_\mathrm{L} = 42$. The video has been converted from cine to mp4 format and trimmed to show two stroke cycles. The first frame of the video represents the start of power stroke for the P5 paddle.

**Video 2.** Example video used for PIV processing for the case of $Re_\mathrm{L} = 365$. The video has been converted from cine to mp4 format and trimmed to show two stroke cycles. The first frame of the video represents the start of power stroke for the P5 paddle.

**Video 3.** Example video used for PIV processing for the case of $Re_\mathrm{L} = 36{,}483$. The video has been converted from cine to mp4 format and trimmed to show two stroke cycles. The first frame of the video represents the start of power stroke for the P5 paddle.

### SUPPLEMENTARY DATA

**SupplementaryData.zip:** compressed (ZIP) folder containing the following comma separated values (CSV) files:
**ReL32_Betas.csv:** Tracked paddle bending angle ($\beta$) versus dimensionless time data for $Re_L = 32$.
**ReL64_Betas.csv:** Tracked paddle bending angle ($\beta$) versus dimensionless time data for $Re_L = 64$.
**ReL27362_Betas.csv:** Tracked paddle bending angle ($\beta$) versus dimensionless time data for $Re_L =$ 27,362.
**ReL54724_Betas.csv:** Tracked paddle bending angle ($\beta$) versus dimensionless time data for $Re_L =$ 54,724.
